# Discovering pathways through ribozyme fitness landscapes using information theoretic quantification of epistasis

**DOI:** 10.1101/2023.05.22.541765

**Authors:** Nathaniel Charest, Yuning Shen, Yei-Chen Lai, Irene A. Chen, Joan-Emma Shea

## Abstract

The identification of catalytic RNAs is typically achieved through primarily experimental means. However, only a small fraction of sequence space can be analyzed even with high-throughput techniques. Methods to extrapolate from a limited data set to predict additional ribozyme sequences, particularly in a human-interpretable fashion, could be useful both for designing new functional RNAs and for generating greater understanding about a ribozyme fitness landscape. Using information theory, we express the effects of epistasis (i.e., deviations from additivity) on a ribozyme. This representation was incorporated into a simple model of the epistatic fitness landscape, which identified potentially exploitable combinations of mutations. We used this model to theoretically predict mutants of high activity for a self-aminoacylating ribozyme, identifying potentially active triple and quadruple mutants beyond the experimental data set of single and double mutants. The predictions were validated experimentally, with nine out of nine sequences being accurately predicted to have high activity. This set of sequences included mutants that form a previously unknown evolutionary ‘bridge’ between two ribozyme families that share a common motif. Individual steps in the method could be examined, understood, and guided by a human, combining interpretability and performance in a simple model to predict ribozyme sequences by extrapolation.

## Introduction

The mapping of genotype to phenotype for functional biopolymers implicitly captures information about structural contacts and mechanism. Fitness landscapes are mathematical maps that relate primary sequence to functional properties, such as catalytic rate enhancement for enzymes or ribozymes. Understanding these landscapes, particularly for RNA, may yield insights into mechanisms as well as the molecular evolution of early life (1,2). The development of quantitative tools and high-throughput experiments for elucidating and analyzing fitness landscapes is therefore a major front in research efforts to understand these systems (3). High-throughput collection of data has been used to characterize the fitness landscapes of RNAs (4–6). For example, kinetic measurement using high-throughput sequencing (e.g., *k*-Seq) is able to measure the activities of tens of thousands of ribozyme sequences (7,8).

Nevertheless, even with high-throughput experimental techniques, only a small fraction of possible sequence space can be sampled, due to both synthetic and analytical limitations. For example, a fully randomized 30-nt region in a ribozyme sequence would yield 10^18^ different sequences, far exceeding current high-throughput sequencing capacity. Therefore, computational methods are required to predict activities for sequences that were not captured by the empirically available data. Such data presents computational challenges for interpretation, and improved analytical techniques are required to quantitatively characterize fitness landscapes and develop models that advance understanding of the genotype-phenotype relationship.

In this work, we focus on an activity that would have been foundational to the genetic code of protein translation, perhaps the greatest evolutionary invention of an early RNA-based prototypical life (9). A key activity of this process is the covalent attachment of amino acids to tRNAs (10), which is catalyzed in contemporary biology by aminoacyl-tRNA protein synthetases. In the pre-protein world, however, this activity may have been achieved by self-aminoacylating ribozymes. This hypothesis is supported by the existence of ribozymes that react with aminoacyl adenylates or other activated substrates (11–14). Prior work (9) determined the catalytic activities of thousands of self-aminoacylating ribozymes that react with 5(4*H*)-oxazolones, considered to be prebiotically relevant substrates (15). Here, we develop and validate a computational method for extracting additional predictive power from the limited experimental data of the fitness landscape.

Existing methods, such as minimum epistasis interpolations (16) and Gaussian processes (17), show promise for interpolating missing data on fitness landscapes and ‘filling in the map’ for regions of sequence space where data is not complete. However, these methods struggle with sparse sampling or require prerequisite knowledge, such as structural data from the Protein Data Bank (18), which are not always accessible for the novel sequences. While modern methods can interpolate fitness landscapes given sufficient sampling, methods for extrapolative predictions looking beyond the boundaries of sampled space are relatively lacking. For example, using information about double mutants of a central sequence to predict activities for triple or quadruple mutants remains an open problem.

A simple extrapolative technique could be based on additivity in the genotype-phenotype map, in which the effects of single mutations on the genotype would be summed to predict the phenotype of the combination. Chemically speaking, additivity corresponds to a separability of chemical moieties that do not interact with one another in the reaction mechanism. For example, a residue that stabilizes the active fold might not interact with a residue that exclusively forms a contact in the transition state. However, additivity is generally not a correct assumption in detail since different residues influence one another through direct contacts or indirect effects (epistasis). Epistatic landscapes feature mutations whose effects are influenced by their genetic context. Attempts to model epistasis include using simple nonlinear functions to capture latent, non-epistatic traits (19–22) and machine learning models (23,24). The former technique performs well for relatively simple systems in which there is a largely additive landscape subject to random variation, but not for more complex landscapes. Machine learning has strong general capability but requires considerable finesse in parameterization and can pose difficulties with interpretation. In one recent study, in silico evolution was performed on ribozyme variants using empirically determined fitness values, with a deep learning perceptron model applied in the final round. While the approach effectively identified neutral mutants of the ribozyme, the perceptron itself constituted a ‘black box’ (25). Therefore, methods that combine performance and interpretability are needed.

One possible approach is to use mathematical language to construct an articulation of epistatic complexity that remains accessible to human insight. The method described here applies information theory to identify regions of sequence space where non-interfering mutations can be exploited to extrapolate beyond the boundaries of the measured space. Instead of fitting a function to the fitness landscape, this method identifies mutations that are likely to yield high activity when combined. Epistasis has previously been analyzed in a probabilistic framework (26). Here we relate the epistatic quantity to information theory and demonstrate its ability to provide a pairwise decomposition of the information contained in the data set. This pairwise representation can be exploited to create a predictive model without fitting parameters.

Noting that mutual information has seen success improving prediction outcomes when integrated into models of the sequence-activity relationship (27–29), we use surprisal (see equation 1, below) and mutual information to calculate a quantity termed ‘epistatic divergence’. We demonstrate that epistatic divergence can be used to derive insights from empirical data that extrapolate beyond the explored regions of their fitness landscapes. Using two families of ribozymes for which the activity of all possible double mutants of a central ‘seed’ sequence had been measured, we predicted and validated points in the sequence space of triple and quadruple mutants with high likelihood of activity. Epistatic divergence identified an evolutionary connection between two ‘islands’ of activity within the fitness landscape, which we validated experimentally. Such extrapolation could be combined with interpolation algorithms to enable greater understanding of fitness landscapes.

This study proposes a representation of interactions in the sequence-activity landscape, in which qualitative properties of a system are articulated mathematically. Representations are an essential part of model development (30,31) that affect the fundamental ability to observe patterns in the data. This epistatic divergence representation explicitly captures the degree to which the sequence-ribozyme activity relationship is epistatic, which subsequently enables precise exploitation of non-interfering mutations for extrapolative predictions.

## Methods

### Construction of epistatic divergence

We construct epistatic divergence in a similar manner to Ostman et al. (26) to compare the degree to which a pair of nucleotide identities affects the activity state versus the degree to which an individual constituent site affects the activity state. The motivation is that a more epistatic nucleotide pair requires the knowledge of both nucleotides jointly to describe the activity state likelihood more accurately, while a less epistatic pair would allow for that description from the individual descriptions of each nucleotide identity. We describe the epistatic divergence using information content (*I*), or surprisal (32). Formally, *I*(*p*(*x*)) is the information content of event *x* with probability *p*(*x*), where

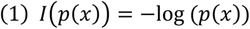

Epistatic divergence is assessed as:

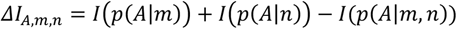

Here, we denote a random variable representing the activity of the ribozyme with *A*. Lower case *m* and *n* specify a genotype at sites M and N. We term a pair of sites relevant to this calculation as a ‘site pair’. Thus, *p*(*A*|*m*) denotes the probability of observing *A* conditioned on genotype *m*, and we have:

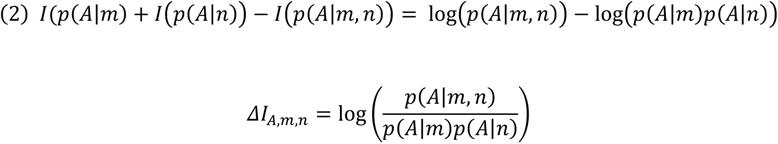

This expresses epistatic interaction using information content. Because we are concerned with pairs of sites (i.e., ‘site pair’, such as 29 and 38), we average over a probability distribution that describes how the various genotypes predict the phenotypes. Thus we use as our distribution *p*(*A*|*m*,*n*):

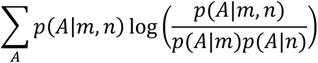

To incorporate information about every possible genotype at a site pair, we sum over all genotypes that are represented in the sample (over the support set of the population). Thus we sum over every combination of activity and genotype states (*A*, *m*, *n*) that has at least one representative in the sample population, as follows:

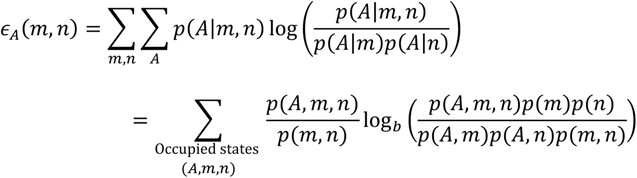

Thus,

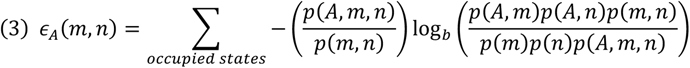

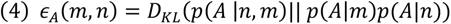

where *D_KL_* is the Kullback-Leibler divergence and *∈_A_* is the epistatic divergence. The quantity *∈_A_* is equal to zero when the interaction between *m* and *n* carries no additional information (i.e., no epistasis, such that the activity of the combined genotype can be completely predicted from the activities of the individual genotypes), or when the sites do not affect activity. We set base *b* = 2, so that information is given in units of bits.

It should be noted that the use of the *D_KL_* is compact notation and does not relate to the use of *D_KL_* to compare two probability distributions, because *p*(*A*|*m*)*p*(*A*|*n*) is not a well-defined probability distribution. Thus, the usage here only results from examining the difference of information contents between informational bodies and weighting it to favor relevance to the empirical distribution, *p*(*A*|*m*,*n*).

Together, the logarithmic terms quantify the degree to which genotypes are statistically dependent within the context of a given genotype, and the weight factor then adjusts the signal such that its intensity depends on the degree to which the genotype explicitly impacts the phenotype. Detailed discussion of the epistatic divergence quantity is provided in the Supplementary Information, Appendices A-C.

### Mutual information to describe the effects of single sites

To assess the single site effect on polymer activity, we use the mutual information, in bits, *N*(*A*;*m*), as follows:

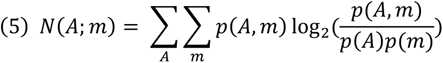

Mutual information is used to interrogate the divergence of the joint state distribution between activity and single-site identity, and the distribution associated with statistical independence, thus quantifying the effect of a given residue on the distribution of the activity classes. In contrast to epistatic divergence, the mutual information assesses single-site contributions rather than interactions between sites with respect to their impact on the activity class.

The activity, *A*, is a continuous value, which we classify into discrete classes of activity by a schema **A** with a set of associated parameters {θ*_i_*}, where *i* indexes the parameters that discretize the activity space. We used two types of classification. The first, used for extrapolative prediction of active sequences, divides sequences into those with activity less than or greater than the activity of a central reference ‘seed’ sequence. The second, used for visualizing the pairwise epistasis contributing fundamental activity, was found by fitting a gaussian curve to normalized activity values. This results in a threshold of four times the baseline activity, that statistically defines whether a sequence can be considered catalytically active or not (33). Median values from experimental replicates were used for the activity metric.

We specify the classification scheme **A**, as follows:

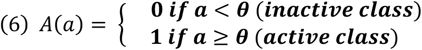

Where *a* is the continuous value of activity and *A* is the discrete class. This representation depends only on a single parameter, θ; however, alternative classification schemes could be defined depending on the needs of a given investigation.

### Extrapolative prediction of active sequences

To predict regions with high-activity ribozymes, we compute ε, epistatic divergence, using the classification threshold θ = *a*_seed_, the activity of the seed sequence at the center of the 2-Hamming distance radius defining the training space. For other characterization, we set θ = *a*_active_, a value determined by fitting normal distributions to the measured background activities of non-catalytic sequences. Ribozyme activities higher than the threshold value were significantly greater than the background activity (i.e., >4 times the background rate) (33).

The epistatic divergence values were plotted to determine which sites were most associated with improvements upon the base activity of the seed sequence. This process identified pairs of nucleotides that are associated with the most epistatic improvements to the activity. Knowledge of these pairs was then combined with insight from mutual information calculations regarding highly informative sites. This two-step process resulted in prediction of non-interfering epistatic pairs, whose genotypes could be combined to extrapolate activity in unexplored regions of sequence space.

### Comparison to established measures of epistasis

We compared the information produced by epistatic divergence with two conventional measures, derived from the additive formulation of epistasis (34). We use two quantities, termed *μ* and *σ*, which are related to the activity of sequences as follows:

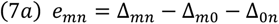

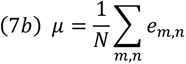

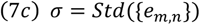

In *μ*, the difference between the change in molecular activity, associated with a double mutant (Δ*_mn_*) and the change described by summing the individual single mutations (Δ*_m_*_0_ and Δ_0*n*_) are averaged over *N* genotypic backgrounds. *σ* is the standard deviation (Std) associated with that set, and describes the variance of effects across different genetic backgrounds. Note that *μ* and *σ* (not ε) are used for denotation of the measures described in equations 7a–7c.

### Experimental data set of ribozyme activities

The data consists of high-throughput (*k*-Seq) activity measurements on two families of ribozymes originally discovered by *in vitro* selection starting from a 21-site variable region. Ribozymes self-aminoacylate by reaction with a tyrosine analog substrate, biotinyl-Tyr(Me)-oxazolone (BYO). Two families, 1B.1 and 1A.1, were chosen for this analysis due to a shared motif. Activity measurements were performed for all sequences within Hamming distance of 2 from the core seed sequences, providing detailed mapping of the localized fitness landscapes. Although no evolutionary pathway had been discovered between these families, the shared motif suggested the possibility of a connection between the two active families. Data from single- and double-mutant sets of these two families were used to produce predictions of high activity within the triple- and quadruple-mutant range of interest, allowing us to explore predictive power beyond the Hamming range of the training set. Data were normalized by background activity, consistent with other work on this data set (33).

### Experimental measurement of ribozyme activity by RT-qPCR

The activities of selected ribozymes were determined by reverse transcription-qPCR assay (RT-qPCR) as previously described (35). DNA sequences were chemically synthesized and polyacrylamide gel electrophoresis (PAGE)-purified by Integrated DNA Technologies. The synthesized DNA sequences were 5′-<u>GATAATACGACTCACTATAGGGAAT</u>GGATCCACATCTACGAATTC-N21-TTCACTGCAGACTTGACGAAGCTG-3′, where the nucleotides upstream of the transcription start site for T7 RNA polymerase are underlined and N21 denotes 21 consecutive nucleotides, which are varied for different ribozyme sequences. The sequences of the N21 region of tested ribozymes are: CCACACTTCAAGCAATCGGTC (S-1B.1-a), CCCCGCTTCAAACAATCGGTC (S1B-29C31G38A), CCCTGCTTCAAACAATCGGTC (S1B-29C30T31G38A), CTGCTTCAAACAATCGGTCTG (S1A-29G), and CTACTTCAAACAATCGGTCTG (S-1A.1-a). RNAs were transcribed using HiScribe T7 polymerase (New England Biolabs) and purified by denaturing PAGE (National Diagnostics). 0.1 μM of RNA samples in the aminoacylation buffer (100 mM HEPES [pH = 7], 100 mM NaCl, 100 mM KCl, 5 mM MgCl_2_, and 5 mM CaCl_2_) were incubated for 90 min with various BYO substrate concentrations (10, 50, 100, 250, 500, and 1000 µM) in the total volume of 100 μL for each sample. The reactions were stopped by removing unreacted substrate using Bio-Spin P-30 Tris desalting columns (Bio-Rad). The RNA concentration of each sample was quantified by Qubit® 3.0 Fluorometer (Thermo Fisher Scientific). To isolate the reacted RNA, streptavidin MagneSphere® paramagnetic beads (Promega) were added to all reacted RNA samples (20 ng RNA for each sample from the dissolved reacted RNA stock solutions) with a volume ratio of 1:1. Samples were incubated for 10 min at room temperature with end-over-end tumbling, followed by three washing steps. The aminoacylated RNAs were eluted with UltraPure™ DEPC-Treated Water (Invitrogen) incubation at 70°C for 1 min. The amounts of aminoacylated RNAs were quantified using iTaq SYBR green mix (#1725150, Bio-Rad) using Bio-Rad^®^ CFX96 Touch^®^ system. The samples were prepared following the manufacturer’s protocol. 2 μL sample were mixed in the total 10 μL RT-qPCR reaction volume with 500 nM of both forward and reverse primers. The forward and reverse primers sequence were 5’-GATAATACGACTCACTATAGGGAATGGATCCACATCTACGA-3’ and 5’-CAGCTTCGTCAAGTCTGCAGTGAA-3’, respectively. A calibration standard curve was measured for each RT-qPCR measurement batch to reduce measurement error. The standard RNA sequence was 5′-GGGAAUGGAUCCACAUCUACGAAUUCAAAAACAAAAACAAAAACAAANUUCACUGCAGACUUGACGAAGCUG-3′ which has the same length (i.e., 71 bps) and primer-complementary regions as the ribozymes used in this study. The standard curve was determined by adding 2 μL standard RNA samples with the concentrations of 1000, 100, 10, 1, and 0.1 pg/μL. Triplicates were performed for each sample. Results were fit to the pseudo-first-order rate equation

*F* = *A*(1 − *e*^-*k*[*BYO*]*t*^), where *F* is the reacted fraction, *A* is the maximum reacted fraction, *t* is the incubation time of 90 min, and *k* is the effective rate constant of the aminoacylation reaction. The two fitting parameters *A* and *k* are poorly estimated individually for low-activity sequences (c.a., *k* < 0.5 min^−1^ · M^−1^), but due to the inverse correlation between estimated *A* and *k* during curve fitting, the product of the estimated *k* and estimated *A* is more accurate (7). Therefore, the product of the two estimated parameters, *kA*, from the pseudo-first-order curve fitting, was used to represent the catalytic activity of ribozymes in the present study.

### Data and code availability

The Python code used in the calculations is available at https://github.com/ncharest/epistatic-divergence. The ribozyme data set is publicly available at the Dryad Digital Repository under DOI 10.25349/D92C9C (https://doi.org/10.25349/D92C9C).

## Results

### Epistatic divergence as a measure of epistasis

We analyzed a data set of ribozyme variants of a self-aminoacylating RNA sequence (9), S-1B.1-a, also referred to as a ‘seed’ sequence here. We compared epistatic divergence against a traditional conception of epistasis, namely the difference from additivity of single mutations. For the conventional measure of epistasis, the average difference (*μ*) of a double mutant’s effect on the activity from the sum of the constituent single mutants was calculated for all pairs (*m*,*n*) (site pairs) (Figure 1A). To determine the effect of a single site mutation, all double mutations applying to that site were included in the averaging, reflecting multiple genetic backgrounds present in the double and single mutant data. The standard deviations (*σ*) of these values was also calculated, indicating the spread of the differences from additivity (Figure 1B). These measures reflect the difference from additivity when considering all possible nucleotide combinations across two sites.

**Figure 1.**
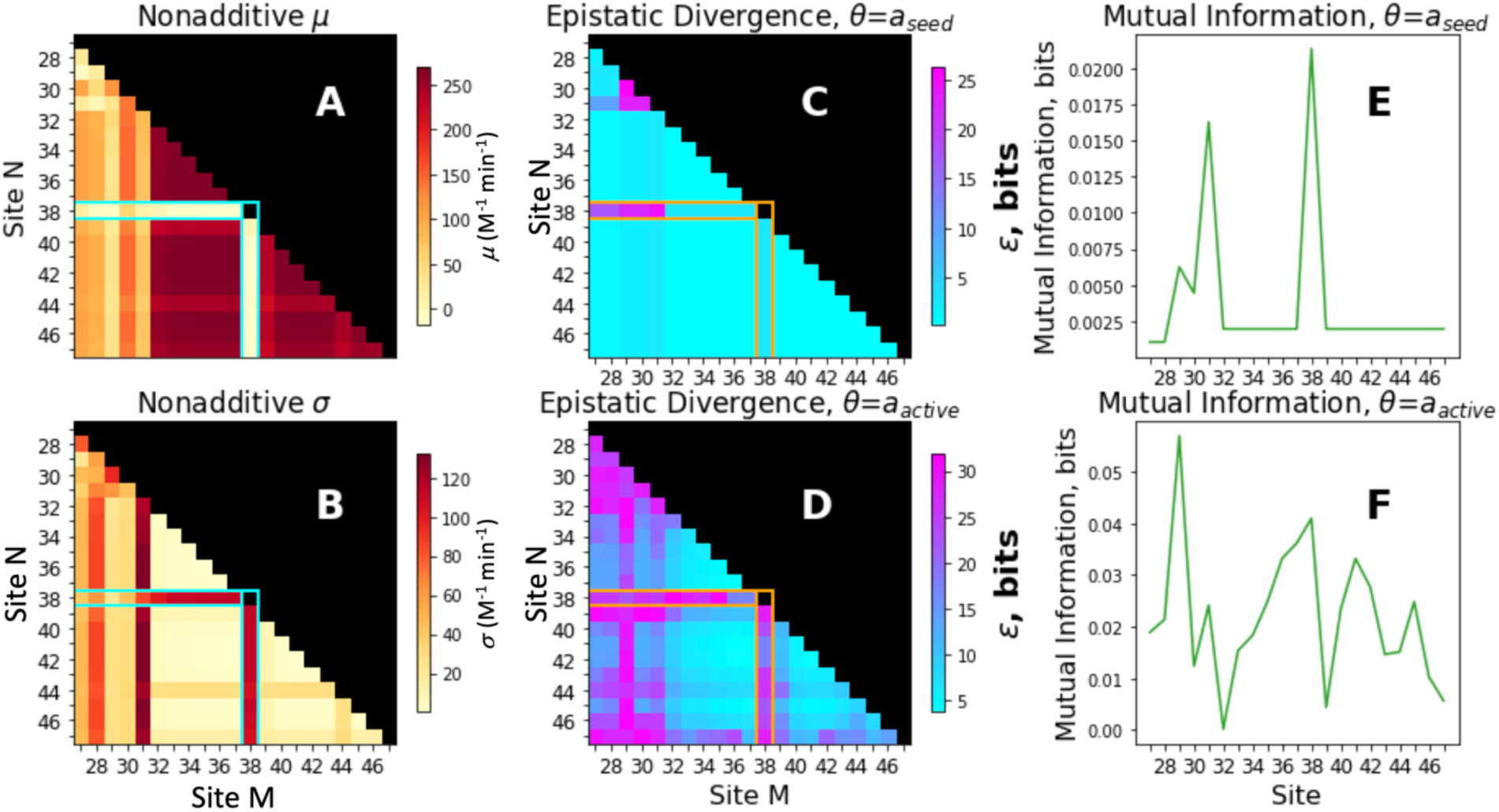
Comparison of epistatic divergence and deviation from additivity for ribozyme S-1B.1-a. (A) The ensemble average of double mutants’ difference from the additive sum of single mutants, a conventional measure of statistical epistasis that shows deviation from additivity. (B) The standard deviations of the calculations from part A. (C) The epistatic divergence computed using the threshold *a_seed_*. The seed sequence defines the sample population; all mutants in the experimental data are a Hamming distance of 1-2 from the seed. The median activity of the seed was taken as *a_seed_*, and any sequence whose median activity was found above *a_seed_* was marked superior while any whose median was found lower was marked inferior. Note that sequences with activity close to the seed sequence may be incorrectly classified due to experimental noise (7). (D) The epistatic divergence calculated using the threshold *a_active_* based on the background catalytic rate determined in prior work (33). (E) The mutual information depicting which sites along the sequence were found to have the greatest relevance to the classification around *a_seed_*. (F) The mutual information calculated around the classification scheme with *a_active_*. The goal of these measures is to detect the site significance to the catalytic activity of the ribozyme.

For epistatic divergence, low values indicate a site pair (*m*,*n*) where the effects of the nucleotide combination were found to be either lacking impact on the activity, or were explainable by considering each site independently, or both. A high value of epistatic divergence indicates a site pair where the combination of nucleotides was found to be important and impactful upon the activity. We calculated the epistatic divergence using two possible values of the classification thresholds θ, namely classified based on activity less than or greater than (or equal to) the seed sequence (θ = *a*_seed_; Figure 1C), or activity above or below the non-catalytic background rate (θ = *a*_active_; Figure 1D). The former threshold (*a*_seed_) is quite stringent (3 out of 63 possible single mutants and 24 out of 1890 possible double mutants (7)) because the seed sequence is a ribozyme that reached high abundance during prior *in vitro* selection (9), indicating high relative activity. This threshold choice was used to focus on sites that may cause activity enhancements close to or greater than the seed sequence. The second threshold (*a*_active_), set at a catalytic rate equal to four times the background (non-catalytic) reaction rate, captures sites that influence whether a sequence is catalytically active at all.

Comparison of the epistatic divergence and the conventional measure of epistasis shows that a site of particular interest is position 38, which exhibits high *σ* despite low *μ*, indicative of highly variable epistasis depending on genetic background. Consistent with this, epistatic divergence is high for site 38, suggesting an important relationship between activity and the nucleotide identity at this site. Importantly, epistatic divergence also highlights other regions of the sequence that do not appear unusual based on the traditional measure of epistasis, particularly when considering highly active sequences (Figure 1C vs 1A, B). Predictions based on the region highlighted by epistatic divergence, but not the traditional measure of epistasis, were tested experimentally (described below). These features demonstrate the ability of epistatic divergence to positively identify regions of interest in the ribozyme.

Conversely, the ability to correctly identify regions that are not of interest for extrapolative combination is also important. An advantage of epistatic divergence in this regard can be seen in the blocks of signal associated with the regions around sites 32-37 along site M and 39-47 on site N (seen as a low signal region in Figure 1D). The *μ* values suggest a consistently large deviation from additivity, while the spreads (*σ*) are quite narrow (Figure 1A, B). These effects are driven by the fact that these locations are, independently, essential to catalytic function. Mutations in these regions essentially eliminate activity, and so any double mutation of them will lead to high *μ* values due to a saturation effect. However, such patterns cannot be taken to reflect true interactions (i.e., mechanistic or structural) in the ribozyme. In contrast, when epistatic divergence is used, these sites are appropriately identified as lacking interactions. Thus, the epistatic divergence measure has high specificity in identifying loci with complex epistatic behavior that might be exploitable, particularly for predicting active sequences beyond the boundary of sequence space in the data set.

### Identifying hotspots of exploitable complexity

To focus on the potential prediction of high-activity sequences, we used the epistatic divergence measure with θ = *a*_seed_ to develop a predictive model. The epistatic divergence highlighted sites where the data showed a particular dependence on pairwise states when considering the distribution of activity (Figure 1C). Such pairwise interactions are expected to be important for high activity of the ribozyme. At the same time, mutual information (between a single site and the activity distribution) identifies single sites that are informative for activity. Combining mutual information and epistatic divergence should therefore identify mutations that are likely to interact synergistically in high activity sequences. We leverage this fact to produce a simple model that maximizes the utility of non-interfering mutations within the landscape.

Specifically, the individual sites 29, 31 and 38 gave the most information about activity, in ascending order (Figure 1E). Epistatic divergence analysis indicated that the combinations of these loci are synergistic. These observations suggested extrapolative predictions for highly active sequences beyond the experimental data.

### Extrapolative prediction of active ribozyme sequences

The epistatic divergence ɛ is the sum of terms describing each represented state, so the contributions from states can be decomposed into contributions from each specific mutant pair and sorted by activity class. We examined the specific epistatic contributions from states containing a combination of the top four individually informative sites (29, 30, 31, and 38) (Figure 2), showing regions where pairwise epistatic effects contained more information than constituent sites considered alone, for the high activity class (using θ = *a*_seed_). Four sites were included in the analysis in order to obtain predictions for quadruple mutants.

**Figure 2.**
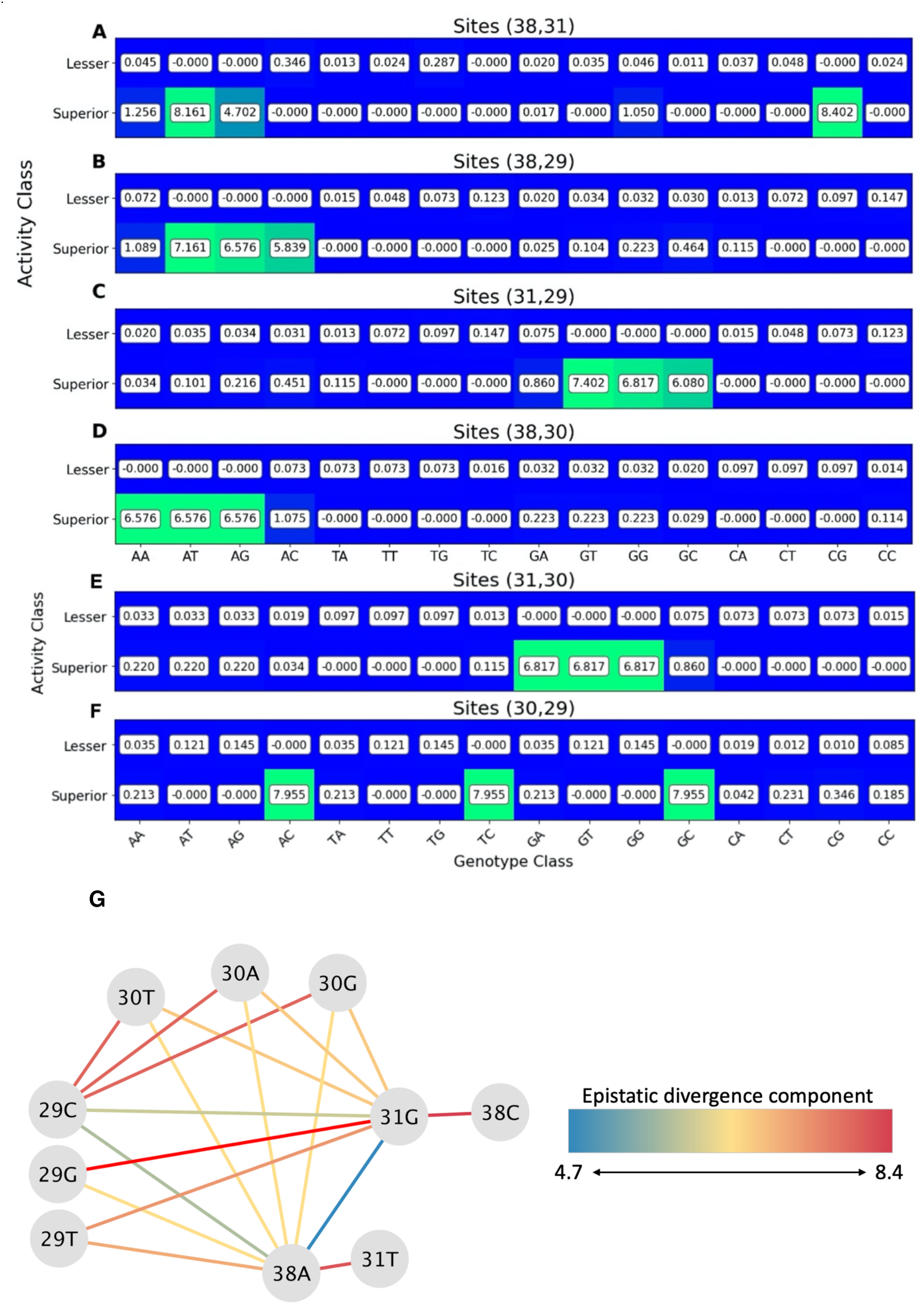
Decomposition of epistatic divergence by sites for ribozyme S-1B.1-a. (A-F) The matrix components of the epistatic divergence calculations for the indicated site pairs. The decomposition was used to identify potentially compatible mutations, which are genotypes associated with improved function over the seed ribozyme that can relate to other pairs of loci. The combinations result in triple or quadruple mutants that are predicted to be likely to exhibit appreciable activity. These predictions extrapolate beyond the mapped fitness landscape. (G) The network of single mutations, in which strong signals are represented by edges. Compatible triple or quadruple mutations are illustrated as completely connected subgraphs. Quantities given are in bits with accompanying heat maps to aid the eye.

In theory, synergistic double mutants might be combined to generate triple and quadruple mutants expected to have high activity. For example, synergistic effects arise from the combinations (38A,31G), (38A,29C), and (31G,29C) (Figure 2A-F). This observation suggested that the new triple mutant (G38A, A31G, A29C) may yield a high activity variant. The following process was used to predict new high activity ribozymes. The epistatic divergence attributed to each site pair (Figure 2A-F) was examined to select potentially informative combinations among the four most highly informative sites (29, 30, 31, and 38). A strong signal was defined as > 4 bits, corresponding to the amount of information needed to completely specify an RNA site pair. This list of strong signals was then searched for compatible combinations that would result in triple or quadruple mutants (Table S1). Two pairs having a common mutation were considered compatible with each other if all of the mutation pairs of the resulting triple mutant were strong signals. For example, (29T,39A) is compatible with (38A,31G) because (29T,31G) is also a strong signal, predicting that the triple mutant (29T,39A,31G) should have high activity. Similarly, two pairs of triple mutants, sharing two of three mutations, were deemed compatible with each other if all pairs of the resulting quadruple mutant were strong signals. For example, (29C,30A,31G), a compatible triple mutant, is compatible with (29C,31G,38A), also a compatible triple mutant, because (30A,38A) is also a strong signal. This procedure can be visualized as a network of single mutations, where nodes (single mutations) are connected if the pair constitutes a strong signal. Compatible triple or quadruple mutants are thus found as completely connected triangles or quadrilaterals (i.e., in which every node is connected to every other node in the subgraph; Figure 2G). This procedure yielded 12 triple mutants and three quadruple mutants that were predicted to have superior activity, assuming that the double mutation information could be combined to produce triple and quadruple mutant predictions. Of these, all of the quadruple mutants were prioritized for experimental testing, since they represent a greater extrapolation compared to triple mutants. Of the triple mutants, half (six) were chosen for experimental testing due to feasibility constraints. The three triple mutants involving the three most informative sites (29, 31, and 38) were all chosen for testing. Of the remainder, triple mutants containing 29C and 38A were prioritized over mutants containing 31G because sites 38 and 29 showed the highest mutual information for θ = *a*_seed_ or θ = *a*_active_, respectively (Figure 1E,F). The sequences selected for testing are given in Table 1.

**Table 1.** Predicted sequences from data on variants of ribozyme S-1B.1-a.

| **Predicted Triple Mutants** | **Predicted Quadruple Mutants** |
| --- | --- |
| s-1B-29C30A38A | s-1B-29C30T31G38A |
| s-1B-29C31G38A | s-1B-29C30A31G38A |
| s-1B-29T31G38A | s-1B-29C30G31G38A |
| s-1B-29C30T38A |  |
| s-1B-29G31G38A |  |
| s-1B-29C30G38A |  |

### Experimental testing of the predicted triple mutant ribozymes

The six triple mutant sequences designed by following the inference process above (Table 1) were used to search the previously obtained high-throughput ribozyme assay data set (33). Although that data set had not been designed to comprehensively cover triple mutants of the seed sequence, some triple mutants had been synthesized by chance along with the variant pool. The distribution of measured activities is shown for analyzable triple mutants from those data (Figure 3A), with an emphasis on triple mutants found to have high activity (>*a*_seed_) (Figure 3B). The six predicted triple mutants indeed had outperformed seed S-1B.1-a, and ranked in the top 30 out of more than 35,000 triple mutants analyzed. Precisions for these measurements are given in Figures S1-S2. Since more active sequences are more likely to have higher relative abundance in the reacted pool (Figure S3), this observation is consistent with the expectation of higher activity level in these triple mutants.

**Figure 3.**
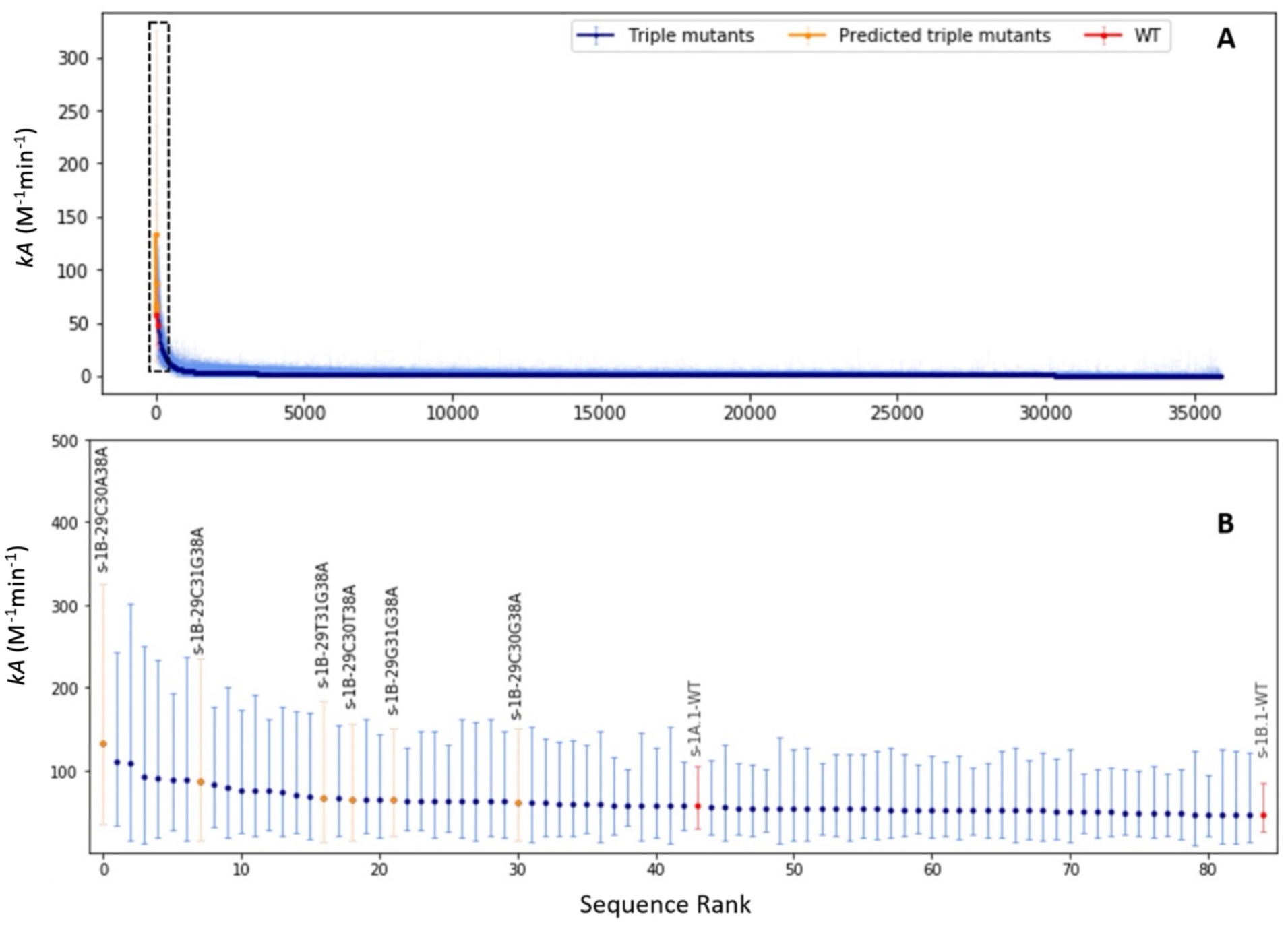
Experimental activity measurements (*k*-Seq (33)) for triple mutants of S-1B.1-a. Measurements are shown as median values with 95% confidence intervals. (A) shows all analyzable triple mutants in the pool, ranked in terms of median *kA* values for activity. (B) shows the top 84 sequences, including seed sequence S-1B.1-a and separate seed sequence, S-1A.1-a (red). The predicted triple mutants (orange) were associated with improvements over S-1B.1-a’s activity and were generally amongst the top-ranking activities. See Figures S1-S3 for measurement precisions.

While the epistatic divergence method shows excellent specificity in identifying high-scoring triple mutants, it should be noted that other triple mutants with top scoring median activities were not detected by the method. This may be due to the fact that the method identifies pairwise contributions to higher activity that are likely compatible, resulting in a set of high order mutants as candidates for testing. The model relies on an expectation that compatible double mutations would not interfere with each other to cause decreased fitness. In other words, in the case of triple mutants, if mutants **AB**, **BC**, and **AC** all have high activity, then mutant **ABC** is predicted to have high activity. (For quadruple mutants, if **AB**, **BC**, **AC**, **AD**, **BD**, and **CD** all have high activity, then **ABCD** is predicted to have high activity.) This expectation is reasonable if epistatic effects diminish at higher orders beyond pairwise interactions (36).

The success of the six predictions suggests this assumption is sometimes appropriate, but violations of this assumption could explain the high-activity triple mutants that were missed in this process.

### Prediction of an evolutionary pathway through a quadruple mutant ribozyme

We also analyzed the epistatic divergence and mutual information for a related ribozyme, S-1A.1-a. Sequence S-1A.1-a and S-1B.1-a are related by a shared motif (Figure 4) offset by 2 sites, but they contain distinct flanking regions and are separated by a total edit distance of 6 (Hamming distance = 16). Interestingly, some mutations suggested by the epistatic divergence analysis of variants of S-1B.1-a were noted to decrease the edit distance to sequence S-1A.1-a. Specifically, A29C, C30T, and G38A would reduce the edit distance between these two ribozyme families. Furthermore, the epistatic divergence analysis for variants of S-1A.1-a (Figure 5) indicated that site 29 is highly informative, and the major signal from epistatic divergence occurs at the (29G, 27A/T/G) pair. Inspection of the sequence alignment indicates that a (29G, 27A) double mutant of S-1A.1-a would reduce the edit distance to sequence S-1B.1-a by two (Figure 4). These considerations indicate a possible connection between the S-1A.1-a and S-1B.1-a families, suggesting there may be an evolutionary path of active ribozyme variants between them.

**Figure 4.**
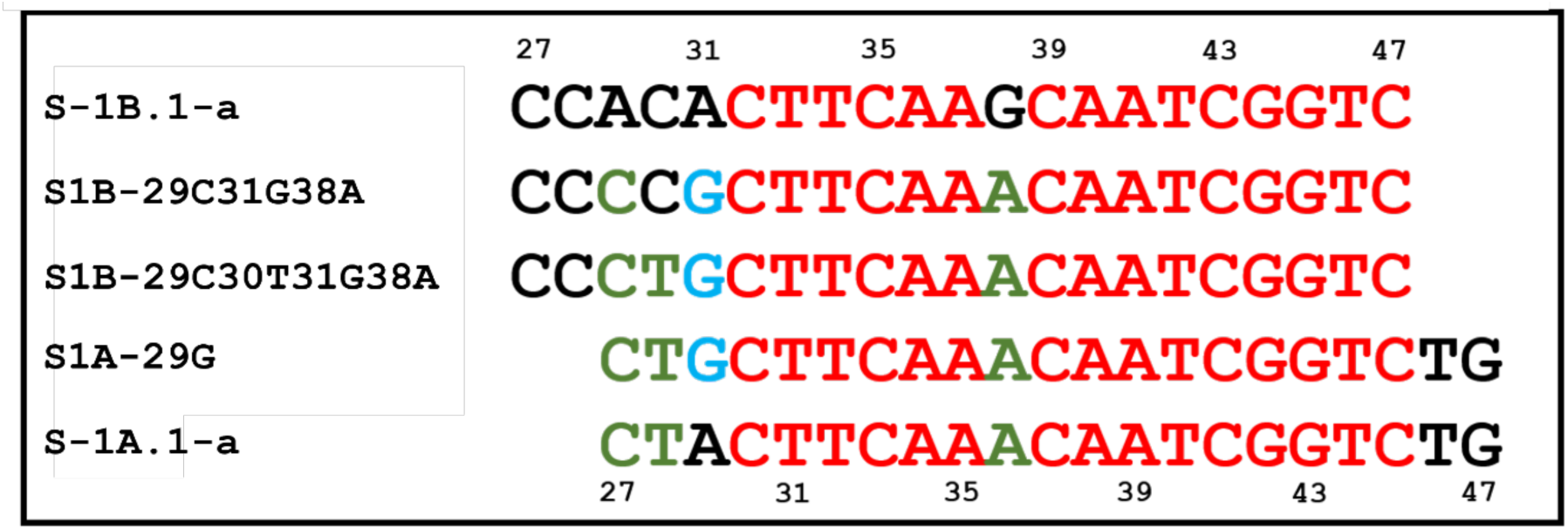
Sequence comparison of seed sequences and mutants indicated by epistatic divergence analysis. In red is the shared motif linking S-1A.1-a and S-1B.1-a families. In green are mutations characteristic of S-1A.1-a and indicated by epistatic divergence analysis as improving S-1B.1-a. In blue are shared residues indicated by epistatic divergence analysis for both S-1A.1-a and S-1B.1-a families. These predictions suggest a possible evolutionary pathway connecting S-1A.1-a and S-1B.1-a.

**Figure 5.**
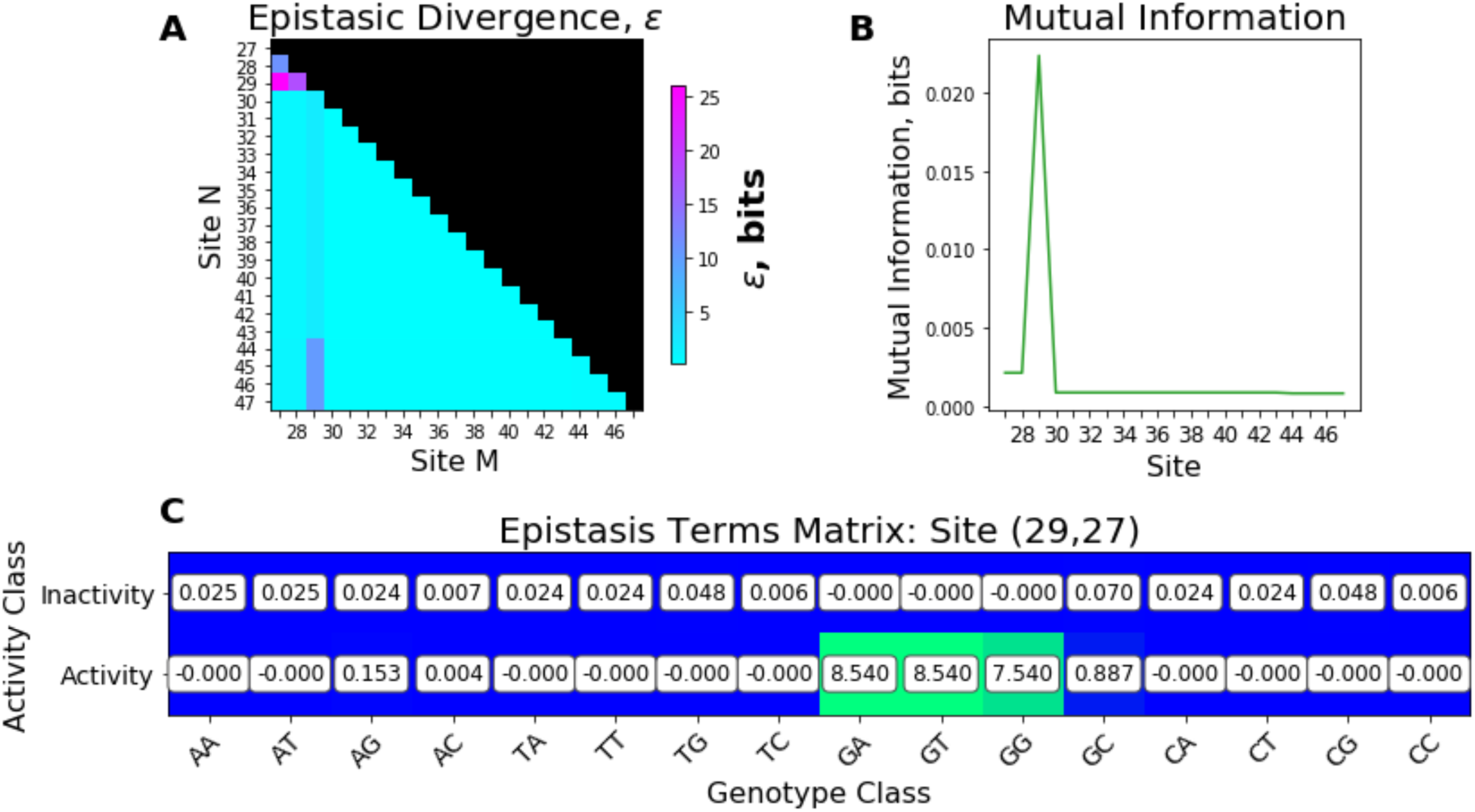
Epistatic divergence analysis for variants of ribozyme S-1A.1-a. Epistatic divergence, mutual information and decomposition from data on single and double mutants of S-1A.1-a. These suggest mutations that bring its sequence closer into alignment with a quadruple mutant of A-1B.1-a that was predicted to be active (S1B-29C30T31G38A).

Thus, epistatic divergence analysis predicted that high activity would occur with mutation of S-1A.1-a to resemble S1B-29C30T31G38A, and conversely that high activity would occur with mutation of S-1B.1-a to resemble S-1A.1-a. This suggested the presence of a specific, a high-activity evolutionary pathway consisting of active mutants to connect these two ribozyme families. We tested the activity of the intermediate mutants (S1A-29G, S1B-29C30T31G38A, and S1B-29C31G38A) individually experimentally. Reaction with the substrate yields a biotinylated product that can be separated using streptavidin beads and quantified by RT-qPCR. Measurement of reaction product over a concentration series allows determination of the catalyzed rate (7). The predicted intermediate mutants indeed exhibited high activity, with some activities being higher than either seed sequence S-1A.1-a or S-1B.1-a, validating the existence of the predicted evolutionary connection (Figure 6).

**Figure 6.**
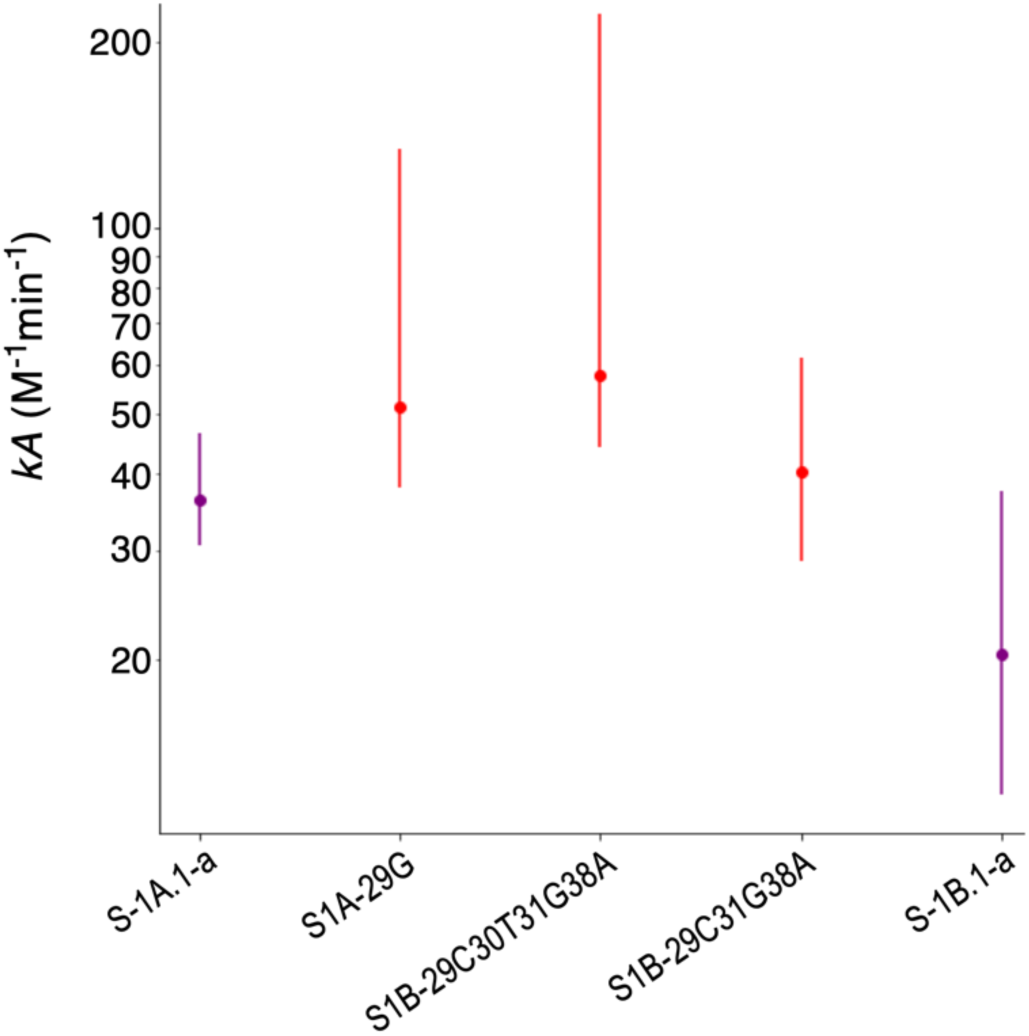
Ribozyme activities along a predicted evolutionary pathway (median and 95% confidence interval), measured by qPCR assay. The endpoints on the x-axis are the seed sequences (purple), with intermediate mutants as shown (red). Mutant sequences were predicted by extrapolation from the data by analyzing epistatic divergence.

## Discussion

The epistatic divergence description of pairwise interactions within the primary sequence of self-aminoacylating ribozymes enabled the extrapolation of previously unknown active sequences, at mutational distances beyond the training space. This was accomplished by considering a data set of activities measured via *k*-Seq experiment, for sequences within two point mutations of a central seed sequence, applying information theory to describe the information content of the distribution of activities in terms of pairs of residue identities, and determining where these pairs possess non-interfering or synergistic behavior that can be exploited to predict highly active sequences beyond the training space. Through this process, the most informative (i.e., highest surprisal) observations of pairwise mutants were used. The most informative observations were combined whenever an internally consistent extrapolation was possible. In other words, epistatic divergence identified where a measurement of activity in double mutants most deviated from the expectation of mutual independence between sites. To generate an extrapolative prediction of triple or quadruple mutants (which were not in the training data set), mutations were combined whenever all of their pairwise interactions were positive (Figure 2G).

Because of the simplicity of this parameter-free approach, the results can be readily interpreted with the language of information theory while simultaneously offering a pragmatic means to identify regions of activity within a fitness landscape, and potential evolutionary pathways, with high specificity.

The epistatic divergence can be mathematically cast as the sum of information contents describing the degree to which a data point contributes to the knowledge of whether a pair of residues predicts an active sample or an inactive sample. Unlike machine learning methods that predict a distribution of activity over the sequences, this method mathematically identifies the parts of the sequence that yield the most information for predicting the activity. This approach systematizes empirical methods that rely on manual curation of significant residues (37). Subsequent analysis then allows the construction of predictions outside the training set, by combining the sequence identities that were associated with the most informative data points within the training set. Mutual information has been previously used for detecting coupled variables in biological contexts. Examples include analyzing combinations of SNPs in genetic studies (29,38) and predicting contacts between residues in protein (39,40) or RNA (41,42) molecules. In this work, we further identify beneficial genotypes using surprisal and combine compatible genotype pairs to detect active higher-order mutants.

Recent work on genotype-phenotype mapping by minimizing epistatic interactions develops a model that allows for maximally locally additive behavior indicated by training observations (16). This assumes an underlying preference for non-epistatic behavior, that is then qualified by the epistasis present in the observations of the training pool. Conversely, the epistatic divergence is not a model in the traditional sense of producing a mathematical construction that can generate predictions. Rather, epistatic divergence is a quantitative representation of how informative a given observation is. In this context, an ‘observation’ is a phenotype class (‘active’ or ‘inactive’) paired with a genotype class (e.g., “A’ in position 38 and ‘G’ in position 29) that is present in the training data set. This defines the support set of the independent variables being used to describe the system. Epistatic divergence quantifies information content relative to the rest of the pool, such that the most informative observations can be used for subsequent prediction building. In our model we combine observations that were identified as both highly informative as well as pertaining to the active class. Because we explicitly compute over pairs, we explicitly capture the interactions, epistatic or otherwise, described by those pairs. This results in predictions that are combinations of the most informative pieces of information in the dataset.

Understanding genotype-phenotype maps, and predicting highly active sequences, are important twin goals in biomolecular engineering. Due to the astronomical size of sequence space, for which the mass of a pool containing every possible protein-coding sequence would readily outstrip the mass of the Earth, computational extrapolation will always be necessary to understand genotype-phenotype maps beyond a few dozen residues. Furthermore, given that the vast majority of sequences are inactive, and that the majority of mutations are deleterious to function, computational methods to make accurate, specific predictions are invaluable for identifying novel functional sequences. In this work, the analysis resulted in prediction of 12 highly active triple mutants (of which 6 were tested experimentally) and 3 highly active quadruple mutants (of which all were tested experimentally). It is notable that 9 out of 9 predicted sequences chosen for testing yielded highly active sequences. In a previous study measuring activities by kinetic sequencing, approximately 5% of all single mutants and 1% of all double mutants were found to have high activity. The previous study using a doped library was not designed to measure all possible triple or quadruple mutants, but many were still measured though at low precision due to a small number of sequencing reads. Of these triple and quadruple mutants, 0.2% or less were found to have high activity (Table S2) (7). Therefore, the epistatic divergence method described here compares favorably in identifying active mutants (9/9) compared with the very low frequency of active mutants from an unbiased sample. While the double mutant data, on which this method is based, was comprehensive (i.e., including all possible double mutants), no data on triple or quadruple mutants was used for the predictions. However, the predictive power of this method is likely to decrease for higher-order mutations, since the method assumes that higher-order epistatic interaction is relatively small when predicting mutants. Progress in increasing the throughput of synthetic and analytical techniques would be useful for building larger experimental data sets to validate predictions.

Furthermore, the pattern of these mutants revealed a previously unknown neutral evolutionary pathway of highly active sequences through the fitness landscape, which joined the two ribozyme families centered on S-1A.1-a and S-1B.1-a. In particular, while the experimental data set used for the analysis here described only the local fitness peaks (within a mutational distance of 2) around sequences S-1A.1-a and S-1B.1-a, the epistatic divergence specifically illuminated multiple high points outside this region, as well as an evolutionary connection that was previously unknown. An experimental approach to the same goal of discovering new fitness peaks and an evolutionary pathway, while not impossible, would have been significantly more laborious.

Machine learning approaches have been applied to the problem of predicting active sequences by extrapolation from mutational data. For example, a random forest model was applied to predict active mutants of a self-cleaving ribozyme (43,44). While often successful in generating predictions, random forest models average over many decision trees and thereby create a difficulty in interpreting the process itself. Deep learning models, such as multilayer perceptrons or Long-Short Term Memory networks (25,44,45), improve the representation of the data and extract features found to be significant to the endpoint being modeled. However, deep models are complex, requiring many parameters, and interpretability remains an unsolved problem (46). In this context, an advantage of the analysis presented here is that the statistical quantities are not based on fitted parameters, but are rather calculated directly from the data, and the prediction process follows well-defined steps from the calculation of epistatic divergence components to the assessment of mutant combinations.

Thus, in the epistatic divergence analysis presented here, the steps of the method are directly interpretable in real terms, and the analysis itself is an interactive process with the data, allowing insight into the genotype-phenotype map. The analysis here shows how epistatic divergence can highlight regions significant to the genotype-phenotype model, and provides means to reliably predict their combinatorial nature from simple, meaningful quantities. This expands the capability to discuss these mappings in rigorous terms and complements the application of more sophisticated modelling methods by offering a method to expose the underlying statistical behaviors. Such mixed-approach analyses are crucial for converting large-scale data sets into specific biochemical knowledge.

## Conclusions

In this work we demonstrated a simple quantity that can be calculated from a large but limited bulk of sequence-activity data to produce a probabilistic representation of the ribozyme fitness landscape. This representation explicitly captures the degree to which a given sequence site possesses epistatic interactions with other sites, enabling precise exploitation of these differing forms of interaction.

Contemporary machine learning efforts frequently rely on the application of “shallow learners” (31), algorithms applied directly to biochemical data with the hope that the sophistication of the algorithm is sufficient to overcome the convolutions obscuring the sequence-activity relationship. However, the choice of representation for the input data significantly impacts not just the interpretability of the model but also the performance of the model (30). With this in mind, the epistatic divergence introduced here is a simple transformation of the data, driven by established information theory. The results are used to develop a simple model that maximizes our extrapolation capabilities, such that we could predict and experimentally validate new points in sequence space having high activity. We demonstrated that epistatic divergence is a sufficient representation to create experimentally relevant extrapolative models using a simple analysis workflow. Future integration of epistatic divergence with sophisticated machine learning algorithms (e.g., (18)) may further improve predictive models of fitness landscapes.

## Supporting information

Supplementary Information for preprint

## Acknowledgements

Nathaniel Charest now works as a Federal Postdoctoral Chemist for the United States Environmental Protection Agency. All work for this manuscript was completed before working for the U.S. EPA. The views expressed in this work are those of the author and do not reflect U.S. EPA opinions or policy.

