## Supplementary Information for preprint for "Discovering pathways through ribozyme fitness landscapes using information theoretic quantification of epistasis"

#### Appendix A

*Analysis of ε, the epistatic divergence*

We can rewrite the expression as follows:

$$\epsilon_{A,m,n} = \sum_{A,m,n} p(A|m,n)[\log\big(p(A|m,n)\big) - \log\big(p(A|m)\big) - \log\,(p(A|n)]$$

$$\epsilon_{A,m,n} = \sum_{A,m,n} p(A|m,n)\left[\log\left(\frac{p(A,m,n)}{p(A,m)p(A,n)}\right) - \log\left(\frac{p(m,n)}{p(m)p(n)}\right)\right]$$

$$= \sum_{A,m,n} p(A|m,n)[\phi_{A,m,n} - \upsilon_{m,n}] = \sum_{A,m,n} p(A|m,n)\phi_{A,m,n} - \sum_{m,n} \upsilon_{m,n}$$

We can inspect each term individually.

*Analysis of Upsilon*

$$\upsilon_{m,n} = \log\left(\frac{p(m,n)}{p(m)p(n)}\right)$$

This term describes the degree to which the sampling of genotypes differs from statistical independence. It corrects epsilon for bias in the sampling of genotypes within the database.

*Analysis of Phi*

$$\phi(A,m,n) = \log\left(\frac{p(A,m,n)}{p(A,m)p(A,n)}\right)$$

It is illustrative to compare this term to another, which we construct as follows:

$$\phi'(A,m,n) = \log\left(\frac{\frac{N[A \cap m \cap n]}{N[A]}}{\left(\frac{N[A \cap m]}{N[A]}\right)\left(\frac{N[A \cap n]}{N[A]}\right)}\right) = \log\left(\frac{p_A(m,n)}{p_A(m)p_A(n)}\right)$$

The operator N[X] denotes the length of set X. ∩ is the intersection operator per convention.

This is a measure of divergence from statistical independence between genotypes m and n within the subset of A

$$\phi'(A,m,n) = \log\left(\frac{N[A]N[A \cap m \cap n]}{N[A \cap m]N[A \cap n]}\right) = \log\left(\frac{N[A \cap m \cap n]}{N[A \cap m]N[A \cap n]}\right) + \log(N[A])$$

We can repeat a similar process for phi.

$$\phi(A,m,n) = \log\left(\frac{p(A,m,n)}{p(A,m)p(A,n)}\right)$$

$$= \log\left(\frac{\frac{N[A\cap m\cap n]}{N_{tot}}}{\left(\frac{N[A\cap m]}{N_{tot}}\right)\left(\frac{N[A\cap n]}{N_{tot}}\right)}\right) = \log\left(\frac{N[A\cap m\cap n]}{N[A\cap m]N[A\cap n]}\right) + \log(N_{tot})$$

Comparing the two we can see the relationship:

$$\phi(A,m,n) - \phi'(A,m,n)$$
$$= \log\left(\frac{N[A\cap m\cap n]}{N[A\cap m]N[A\cap n]}\right) + \log(N_{tot}) - \log\left(\frac{N[A\cap m\cap n]}{N[A\cap m]N[A\cap n]}\right)$$
$$- \log(N[A]) = -\log\,(p(A))$$

$$\phi(A,m,n) = \phi'(A,m,n) - \log\big(p(A)\big) = \phi'(A,m,n) + I\big(p(A)\big)$$

Where I(A) is the self-information of p(A).

*Analyzing Terms Together*

Thus, we can write:

$$\epsilon_{A,m,n} = \sum_{A,m,n} p(A|m,n)[\phi'_{A,m,n} + I(A) - \upsilon_{m,n}]$$

This form allows us to describe each contribution intuitively.

$\phi'$, as derived, is the divergence of genotype pairs from statistical independence within the activity set. This contribution can be zero, positive or negative depending on whether the approximated distribution of joint state m,n within activity set A exceeds the statistically independent expected representation or under performs. The next term, I(A), represents the information gained by simply measuring the observed activity class with the genotype at all. A rarer activity class will be more informative when observed with a given genotype, so this is factored into the general measure of how informative observing the site pair is. Finally, υ corrects for the measured imbalance of genotype sampling. If there are simply more samples of a an m,n pair in the data pool than what would be expected from statistical independence, then the information in $\phi'$ requires correction so that it better represents the distribution divergence due to examining the A subset.

Together these terms quantify how informative a given genotype-phenotype observation is when considering to what degree the genotype is allowing us to predict the phenotype. To account for all genotypes m,n and all phenotypes to provide a grand measure for the site number pair, we sum over all represented states (the support set) of our population.

*The Weighting Term p(A|m,n)*

The $p(A|m,n)$ allows us to interpret epistatic divergence as an expected amount of information emitted from a site pair regarding the degree of conditioning of phenotype upon the genotype at the site loci using the available observation pool.

### Appendix B: Figures Of $\phi'$

This appendix provides figures showing the calculation of epsilon with only $\phi' - \upsilon$ for the sum kernel. It demonstrates that for this observation pool the observation of the active, rare class with a genotype tends to dominate the computation, whereas the approximated divergence from statistical independence of genotypes within the activity classes played a relative minor role. μ and σ are given with units $M^{-1}$ $min^{-1}$.

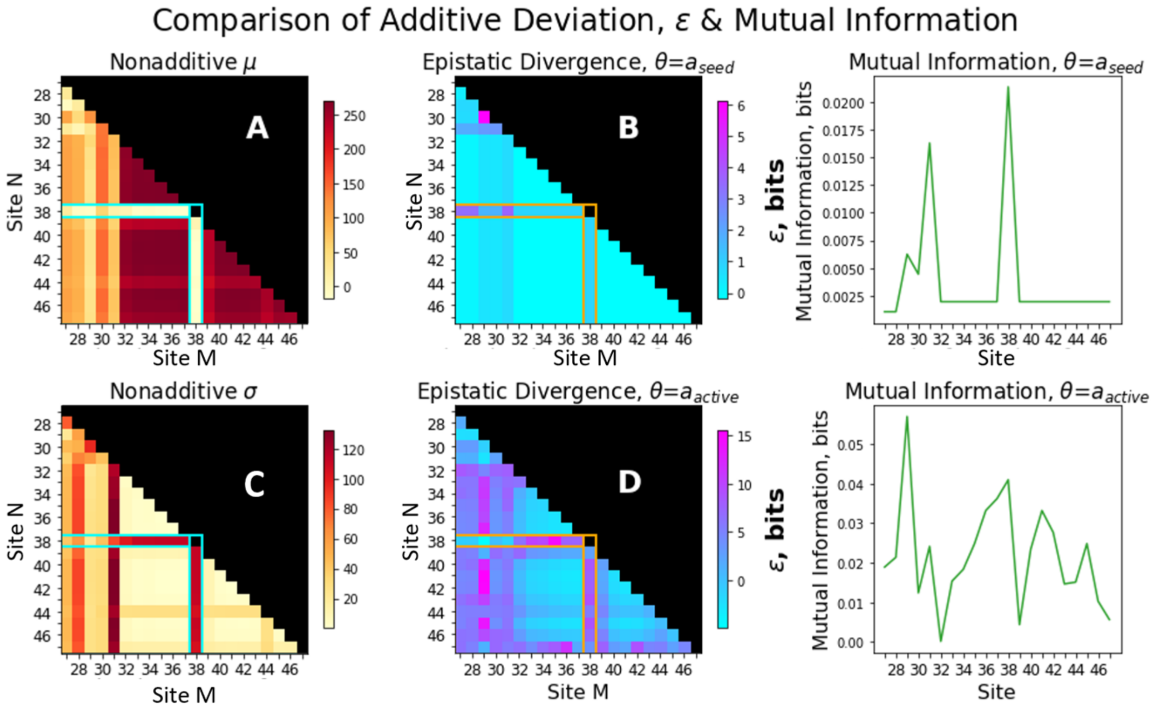


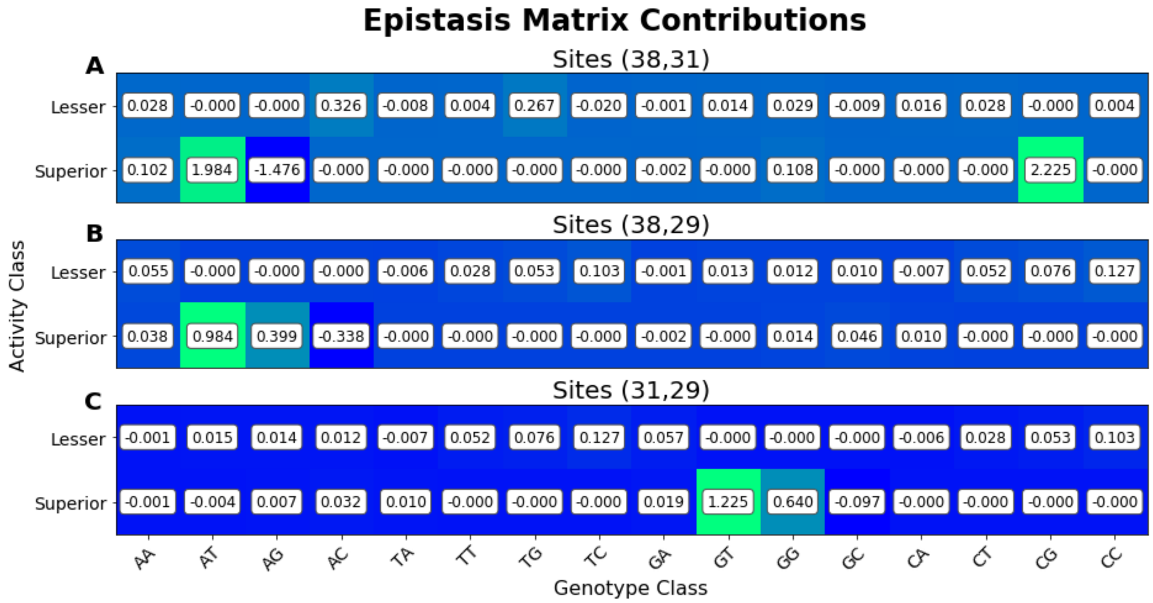

### Appendix C: Toy Models

These are two site models providing examples on how to calculate the epistatic divergence and provide some intuition around it.

*Example 1: Trivial Case*

| **m/n** | **A** | **T** | **C** | **G** |
|---|---|---|---|---|
| **A** | **0** | **0** | **0** | **0** |
| **T** | **0** | **0** | **0** | **0** |
| **C** | **0** | **0** | **0** | **0** |
| **G** | **0** | **0** | **0** | **0** |

1 distinct state:

$p(A|m,n) = 1.0$

$p(A| m) = 1.0$

$p(A| n ) = 1.0$

$\log(p(A|m,n)) - \log(p(A|m)) - \log(p(A|n)) = 0.0$

The null case is a trivial example showing conditions in which there is no epistatic divergence, as no value of m affects the behavior of A and no value of n affects the value of A. Thus, m and n are completely decoupled.

*Example 2: AND*

| **M/N** | **A** | **T** | **C** | **G** |
|---|---|---|---|---|
| **A** | **1** | **0** | **0** | **0** |
| **T** | **0** | **0** | **0** | **0** |
| **C** | **0** | **0** | **0** | **0** |
| **G** | **0** | **0** | **0** | **0** |

3 distinct states:

$p_1(A|m,n) = p_2(A|m,n) = p_3(A|m,n) = 1.0$

$p_1(A|m) = 0.25$, $p_1(A|n) = 1.0$ {note symmetry and arbitrary m,n labels}

$p_2(A|m) = 0.75$, $p_2(A|n) = 0.75$

$p_3(A|m) = 1.0$, $p_3(A|n) = 1.0$

--------

$State_1 = \log(1.0) - \log(0.25) - \log(0.25) = 4$

$State_2 = 0.415$

$State_3 = 0.0$

$1*State_1 = 4$

$6*State_2 = 2.49$

$9*State_3 = 0.0$

---

Epistatic Divergence = 4 + 2.49 + 0.0 = 6.49

This system has epistatic behavior in the form of requiring an A,A genotype for activity, however when considering the behavior of the inactive class there is relatively low complexity. Thus, the value is not as high as one might initially expect.

Compare this result to the base pair model, in which both active and inactive states show epistatic behavior.

*Example 3: Inclusive OR*

| **M/N** | **A** | **T** | **C** | **G** |
|---|---|---|---|---|
| **A** | **1** | **1** | **1** | **1** |
| **T** | **1** | **0** | **0** | **0** |
| **C** | **1** | **0** | **0** | **0** |
| **G** | **1** | **0** | **0** | **0** |

3 distinct states:

$p_1(A|m,n) = p_2(A|m,n) = p_3(A|m,n) = 1.0$

$p_1(A|m) = 0.25$, $p_1(A|n) = 1.0$ {note symmetry and arbitrary m,n labels}

$p_2(A|m) = 0.75$, $p_2(A|n) = 0.75$

$p_3(A|m) = 1.0$, $p_3(A|n) = 1.0$

--------

$State_1 = \log(1.0) - \log(0.25) - \log(1.0) = 2$

$State_2 = 0.830$

$State_3 = 0.0$

$6 * State_1 = 12$

$9 * State_2 = 7.47$

$1 * State_3 = 0.0$

---

Epistatic Divergence = 12 + 7.47 + 0.0 = 19.47

This value is reasonably high, which may be intuitively at odds if one sees the behavior of the genotypes as disconnected, and the activity class only depending on whether a single gene has the correct value.

The interaction comes from the fact that if one gene is 'correct', then the other gene is free to take whichever value it wants without affecting the active state. This is a fairly significant interaction, and thus is registered within the computation.

*Example 4: Exclusive OR*

| **M/N** | **A** | **T** | **C** | **G** |
|---|---|---|---|---|
| **A** | **0** | **1** | **1** | **1** |
| **T** | **1** | **0** | **0** | **0** |
| **C** | **1** | **0** | **0** | **0** |
| **G** | **1** | **0** | **0** | **0** |

3 distinct states:

$p_1(A|m,n) = p_2(A|m,n) = p_3(A|m,n) = 1.0$

$p_1(A|m) = 0.25$, $p_1(A|n) = 0.25$ {note symmetry and arbitrary m,n labels}

$p_2(A|m) = 0.75$, $p_2(A|n) = 0.25$

$p_3(A|m) = 0.25$, $p_3(A|n) = 0.75$

--------

$State_1 = \log(1.0) - \log(0.25) - \log(0.25) = 4$

$State_2 = 2.415$

$State_3 = 0.830$

$1*State_1 = 4$

$6*State_2 = 14.49$

$9*State_3 = 7.47$

---

Epistatic Divergence = 4 + 14.49 + 7.47 = 25.96

The interaction here is like the prior case of inclusive OR. The value is higher because now the activity state is affected if both genotypes are 'correct', which is a more complex interaction.

*Example 5: Base Pairing*

| M/N | A | T | C | G |
|---|---|---|---|---|
| **A** | **0** | **1** | **0** | **0** |
| **T** | **1** | **0** | **0** | **0** |
| **C** | **0** | **0** | **0** | **1** |
| **G** | **0** | **0** | **1** | **0** |

3 distinct states:

$p_1(A|m,n) = p_2(A|m,n) = p_3(A|m,n) = 1.0$

$p_1(A|m) = 0.25$, $p_1(A|n) = 0.25$ {note symmetry and arbitrary m,n labels}

$p_2(A|m) = 0.75$, $p_2(A|n) = 0.75$

--------

$State_1 = \log(1.0) - \log(0.25) - \log(0.25) = 4$

$State_2 = 0.83$

$4*State_1 = 16$

$12*State_2 = 9.96$

---

Epistatic Divergence = 9.96 + 16 = 25.96

A high epistatic divergence that is consistent with the fact activity class depends entirely on the relationship between the genotypes.

**Table S1. Strong signal pairs (edges) in epistatic divergence decomposition.** Predicted mutants (completely connected subgraphs) are listed. Those that were selected for analysis are highlighted in green. Note that the cutoff value for a "strong" signal is a parameter of this analysis. We used a cutoff of at least 4 bits, representing the amount of information that would be used to completely specify a given RNA site pair. However, other thresholds may be useful for other analyses.

| **Edges** | |
|---|---|
| 38A | 31T |
| 38A | 31G |
| 38C | 31G |
| 38A | 29T |
| 38A | 29G |
| 38A | 29C |
| 31G | 29T |
| 31G | 29G |
| 31G | 29C |
| 38A | 30A |
| 38A | 30T |
| 38A | 30G |
| 38A | 30A |
| 38A | 30T |
| 38A | 30G |
| 31G | 30A |
| 31G | 30T |
| 31G | 30G |
| 30A | 29C |
| 30T | 29C |
| 30G | 29C |

| **Completely connected subgraphs** | | | | ***type*** |
|---|---|---|---|---|
| 29C | 30G | 31G | 38A | *quad* |
| 29C | 30T | 31G | 38A | *quad* |
| 29C | 30A | 31G | 38A | *quad* |
| 29T | | 31G | 38A | *triple* |
| 29G | | 31G | 38A | *triple* |
| | 30G | 31G | 38A | *triple* |
| | 30A | 31G | 38A | *triple* |
| | 30T | 31G | 38A | *triple* |
| 29C | 30T | | 38A | *triple* |
| 29C | 30A | | 38A | *triple* |
| 29C | 30G | | 38A | *triple* |
| 29C | | 31G | 38A | *triple* |
| 29C | 30T | 31G | | *triple* |
| 29C | 30A | 31G | | *triple* |
| 29C | 30G | 31G | | *triple* |

**Figure S1. Distribution of *kA* Precisions.** Precision for *kA* measured in the *k*-Seq experiment was evaluated using fold-range from bootstrapping (i.e., the ratio between the 97.5 percentile value and 2.5 percentile value). The number of triple mutants with different precision is shown as a histogram (left y-axis) and the accumulated fraction of mutants within different levels of precision is shown as a stepwise curve (right y-axis). Most of the triple mutants are not precisely estimated (e.g., fold range > 10) due to the limit of sequencing depth.

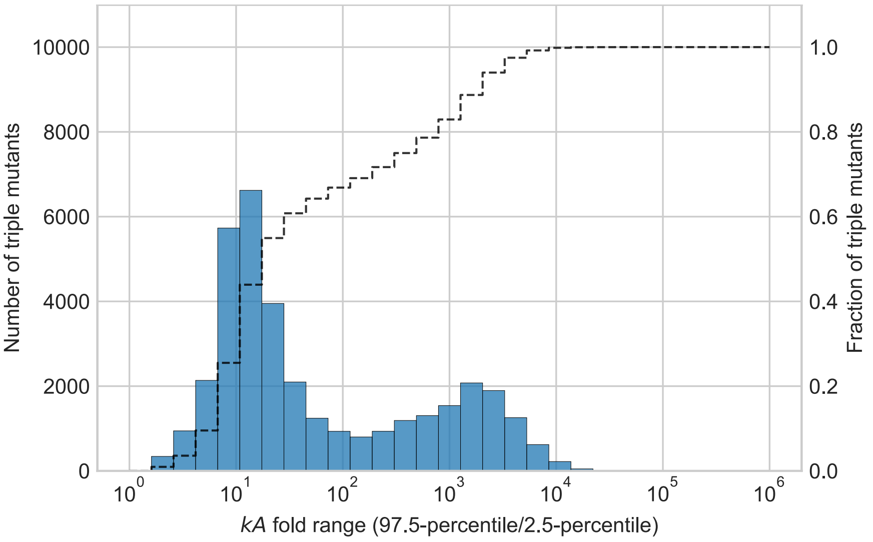

**Figure S2. Distribution of *kA* Precision for Top 80 Median Activities.** Number of mutants vs. precision for *kA* measured in the *k*-Seq experiment, for triple mutants with top 80 median activities. Mutants with higher activities are more precisely estimated in that almost all 80 mutants have fold-range < 10.

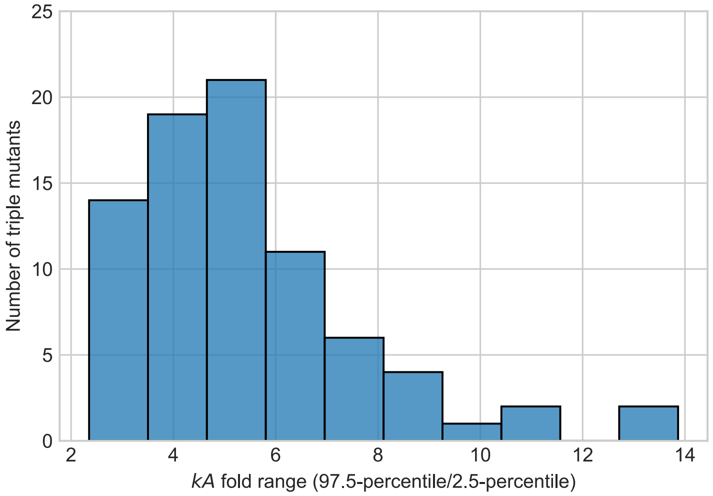

**Figure S3. Scatter Plot of Median *kA* vs. *kA* Precisions.** The distribution of the activity (median *kA*) and measurement precision (fold range for *kA*) for all triple mutants analyzed in the *k*-Seq experiments (blue circles) vs. triple mutants predicted from the model (orange stars). Mutants with lower estimated activities (e.g., < 10 $min^{-1} \cdot M^{-1}$) have wide range of precisions while those with higher estimated activities (e.g., > 10 $min^{-1} \cdot M^{-1}$), including predicted triple mutants, can be precisely estimated (fold range < 10).

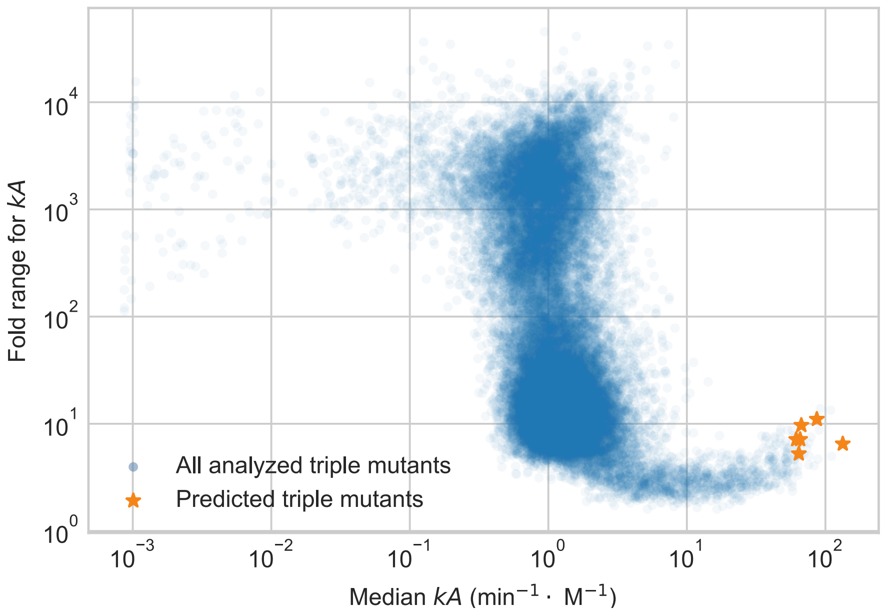

**Table S2.** Number of mutant sequences having activity greater than or equal to the seed sequence. From Shen et al., 2021 (Reference 7).

| Number of mutations | Number of possible unique sequences ($N_u$) | Number of sequences measured by *k*-Seq ($N_m$) | Coverage ($N_m/N_u$) | Number of sequences with activity ≥ $a_{seed}$ as measured by *k*-Seq ($N_a$) | Fraction of high activity sequences ($N_a/N_m$) |
|---|---|---|---|---|---|
| 0 | 1 | 1 | 1 | 1 | 1 |
| 1 | 63 | 63 | 1 | 3 | 0.048 |
| 2 | 1890 | 1890 | 1 | 24 | 0.013 |
| 3 | 35910 | 35907 | ~1.000 | 83 | 0.002 |
| 4 | 484785 | 162592 | ~0.335 | 206 | <0.001 |
